# Preserved wake-dependent cortical excitability dynamics predict cognitive fitness beyond age-related brain alterations

**DOI:** 10.1101/644096

**Authors:** Maxime Van Egroo, Justinas Narbutas, Daphne Chylinski, Pamela Villar González, Pouya Ghaemmaghami, Vincenzo Muto, Christina Schmidt, Giulia Gaggioni, Gabriel Besson, Xavier Pépin, Elif Tezel, Davide Marzoli, Caroline Le Goff, Etienne Cavalier, André Luxen, Eric Salmon, Pierre Maquet, Mohamed A. Bahri, Christophe Phillips, Christine Bastin, Fabienne Collette, Gilles Vandewalle

## Abstract

Age-related cognitive decline arises from alterations in brain integrity as well as in sleep-wake regulation. Here, we investigated whether preserved sleep-wake regulation of cortical function during wakefulness could represent a positive factor for cognitive fitness in aging, independently of early age-related changes in brain structure. We quantified cortical excitability dynamics during prolonged wakefulness as a sensitive marker of age-related alteration in sleep-wake regulation in 60 healthy older individuals (50-69y; 42 women). Brain structural integrity was assessed with amyloid-beta- and tau-PET, and with MRI. Participants’ cognition was extensively investigated. We show that individuals with preserved cortical excitability dynamics exhibit better cognitive performance, particularly in the executive domain, which is essential to successful cognitive aging. Critically, this association remained significant after accounting for brain integrity measures. Preserved dynamics of basic brain function during wakefulness could be essential to cognitive fitness in aging, independently from age-related brain structural modifications that can ultimately lead to dementia.

## Introduction

Aging is associated with an overall cognitive decline triggered in part by a progressive degradation of brain structure. Limited but significant neuronal loss takes place during healthy adulthood [1]. In addition, tau protein, which stabilizes axonal structure, and amyloid-beta (Aβ) protein, a peptide directly related to neuronal activity, progressively aggregate in the brain over the lifespan to form neurofibrillary tangles (NFTs) and senile plaques, respectively [2]. Tau NFTs, Aβ plaques, and neurodegeneration favor cognitive decline [3]. They are considered as major underlying causes of dementia and constitute the hallmarks of Alzheimer’s disease (AD) [4]. However, age-related changes in brain structure go undetected for decades: tau protein aggregation takes place as early as during the second decade of life in the brainstem, while Aβ aggregates can be detected around the 4^th^ decade in the neocortex [5].

One of the first signs of AD- and age-related brain integrity degradation may reside in alterations in the regulation of sleep and wakefulness [6]. Sleep-wake disruption is indeed strongly associated with AD neuropathology [7]: grey matter (GM) integrity has been associated with measures of sleep quality, including sleep slow waves characteristics, in cross-sectional and longitudinal studies [8,9]. Aβ and tau burdens in healthy older individuals have been associated with the amount of slow waves generated during non-rapid eye movement (NREM) sleep [10,11]. Importantly, Aβ burden has been reported to affect memory performance through its impact on sleep slow waves in elderly individuals (~75 y) [11]. In addition, the presence of preclinical Aβ plaque pathology, assessed through PET imaging or cerebrospinal fluid collection, is associated with fragmentation of the entire rest-activity cycle, i.e. encompassing both sleep and wakefulness [12]. Whether age-related alteration in brain structure may affect sleep-wake regulation of daytime brain activity is unknown, however.

Sleep and wakefulness are regulated by two fundamental processes: sleep homeostasis, which keeps track of time awake, and circadian rhythmicity, which temporally organizes physiology and behavior [13,14]. The strength of both processes seems to decrease with age, resulting in dampened dynamics of sleep-wake rhythms and reduced variations in brain activity both during sleep and prolonged wakefulness [13]. The generation of slow waves during sleep, which is associated with the dissipation of sleep need, is reduced in aging [15]. Likewise, cortical excitability [16], a basic aspect of brain function implicated in age-related cognitive decline [17,18], shows less variations during prolonged wakefulness. Age-related alterations in the regulation of sleep and wakefulness are not only associated with current cognition [19,20], but also predicts future cognitive trajectories, including the risk of developing dementia [12,21–23]. Importantly, some of the changes in sleep-wake regulation take place as early as in middle-aged individuals (> 40y) [15]. Whether the early alterations in sleep and wakefulness regulation and their potential cognitive consequences are systematically related to age-related alterations in brain structure remains unknown.

The goals of the present study were threefold. First, we assessed whether sleep-wake regulation of brain function during wakefulness is linked to age-related alterations in brain structure in relatively young and healthy older individuals (50-70 years). We hypothesized that the dynamics of cortical excitability during wakefulness would be related to both Aβ and tau burden, taking into account any potential neurodegeneration. We further investigated whether sleep-wake regulation of brain activity during wakefulness is associated with cognitive fitness. Based on previous findings [16], we anticipated that cortical excitability dynamics would be associated with executive performance. Finally we tested whether these putative links would be independent of Aβ and tau burden as well as neurodegeneration. We postulated that the inclusion of the three markers of brain integrity in our statistical models would at least decrease, if not remove, the association between cortical excitability during wakefulness and cognition.

## Results

In a multi-modal cross-sectional study (**Figure 1A**), 60 healthy and cognitively normal late middle-aged individuals (42 women; age range 50-69 years, mean ± SD = 59.6 ± 5.5 years; **Table 1**) underwent structural MRI to measure GM volume, as well as ^[18F]^Flutemetamol and ^[18F]^THK-5351 PET imaging to quantify Aβ and tau burden, respectively. Participants’ cognitive performance while well-rested was assessed with an extensive neuropsychological task battery probing memory, attention, and executive functions. After a week of regular sleep-wake schedule, participants’ habitual sleep was recorded in-lab under EEG to quantify slow waves generation during non-rapid eye movement (NREM) sleep. A wake-extension protocol started on the following day and consisted of 20h of continuous wakefulness to trigger a moderate realistic sleep-wake challenge under strictly controlled constant routine conditions [14]. Cortical excitability over the frontal cortex was measured 5 times over the 20h protocol, using transcranial magnetic stimulation combined with an electroencephalogram (TMS-EEG) (**Figure 1B-D**) [16,24]. Mean time between wake-extension protocol and cognitive assessment was 30.6 ± 37.7 days. Mean time between wake-extension protocol and brain integrity assessments was 56.7 ± 78.9 days for MRI, 121.3 ± 97.1 days for Aβ-PET, and 116.3 ± 108.3 days for Tau-PET.

**Table 1.** Sample characteristics (mean ± SD).

|  | **n = 60** |
| --- | --- |
| Sex | 42w/18m |
| Age | 59.6 ± 5.5 |
| Education | 15.4 ± 3.2 |
| Right-handed | 59 |
| Ethnicity | Caucasian |
| Dementia Rating Scale | 142.1 ± 2.3 |
| Raven's Progressive Matrices | 50.5 ± 4.9 |
| Mill Hill Vocabulary Scale | 26.9 ± 3.9 |
| BMI (kg/m<sup>2</sup>) | 24.6 ± 2.9 |
| Anxiety | 2.6 ± 2.5 |
| Mood | 4.4 ± 4.7 |
| Caffeine (cups/day) | 3.6 ± 1.9 |
| Alcohol (doses/week) | 3.9 ± 4.0 |
| Sleep quality | 5.1 ± 3.0 |
| Daytime sleepiness | 6.2 ± 3.9 |
| Chronotype | 53.8 ± 8.3 |
| Clock time of dim light melatonin onset (hh:min, PM) | 08:20 ± 00:59 |
| In-lab baseline sleep duration (min, EEG) | 388.0 ± 44.3 |
| In-lab baseline sleep efficiency, including N1 stage (% , EEG) | 82.6 ± 9.6 |
| Baseline sleep time (hh:min, PM) | 10:47 ± 00:36 |
| Baseline wake time (hh:min, AM) | 06:46 ± 00:43 |
| Grey matter volume (% of total volume) | 0.41 ± 0.04 |
| [<sup>18</sup>F]Flutemetamol (SUVR) | 1.16 ± 0.08 |
| [<sup>18</sup>F]THK-5351 (SUVR) | 1.32 ± 0.10 |

- 1 Anxiety was measured by the 21-item Beck Anxiety Inventory [51]; mood by the 21-item Beck - 2 Depression Inventory II [52]; caffeine and alcohol consumption by self-reported questionnaires; - 3 sleep quality by the Pittsburgh Sleep Quality Index [53]; daytime sleepiness by the Epworth - 4 Sleepiness Scale [54]; chronotype by the Horne-Östberg questionnaire [55].

**Figure 1.**
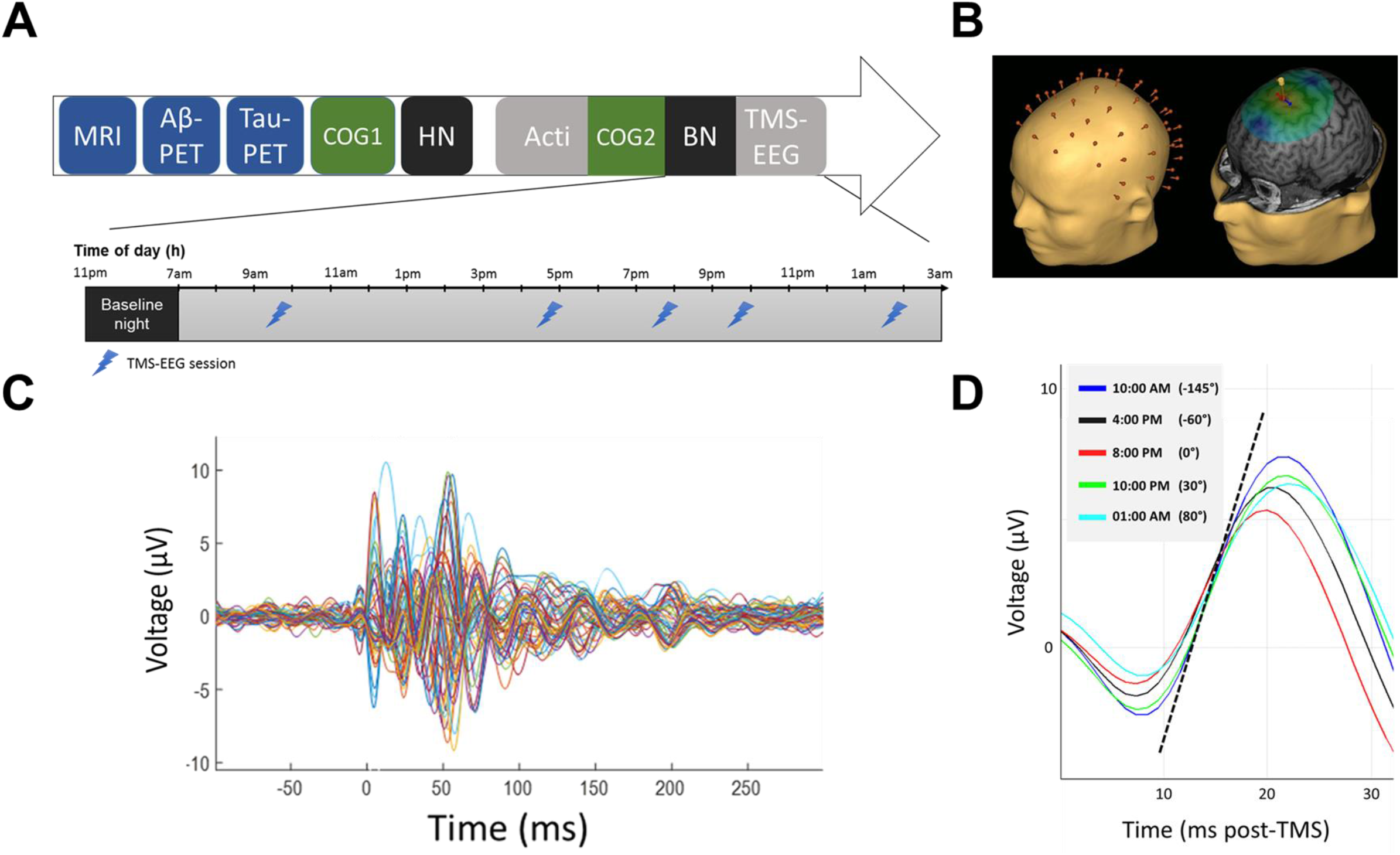
Study design and cortical excitability assessment. **A)** Overview of the whole experimental protocol for a representative subject with bedtime at 11:00PM and wake time at 07:00AM; HN: habituation night with polysomnography; BN: baseline night under EEG recording. **B)** Cortical excitability over the frontal cortex was assessed using neuronavigation-based TMS coupled to EEG. *Left*: reconstructed head with electrodes position; *Right*: representative location of TMS coil and stimulation hotspot with electric field orientation. **C)** Butterfly plot of TMS-evoked EEG response over the 60 electrodes (−100 ms pre-TMS to 300 ms post-TMS; average of ~250 trials). **D)** Representative TMS-response (0-32 ms post-TMS) in the 5 TMS-EEG sessions with indicative clock time and circadian phase (15° = 1h). Cortical excitability was computed as the slope (μV/ms) of the first component of the TMS-response at the electrode closest to the hotspot (dotted line: example for 10:00AM session).

**Figure 2.**
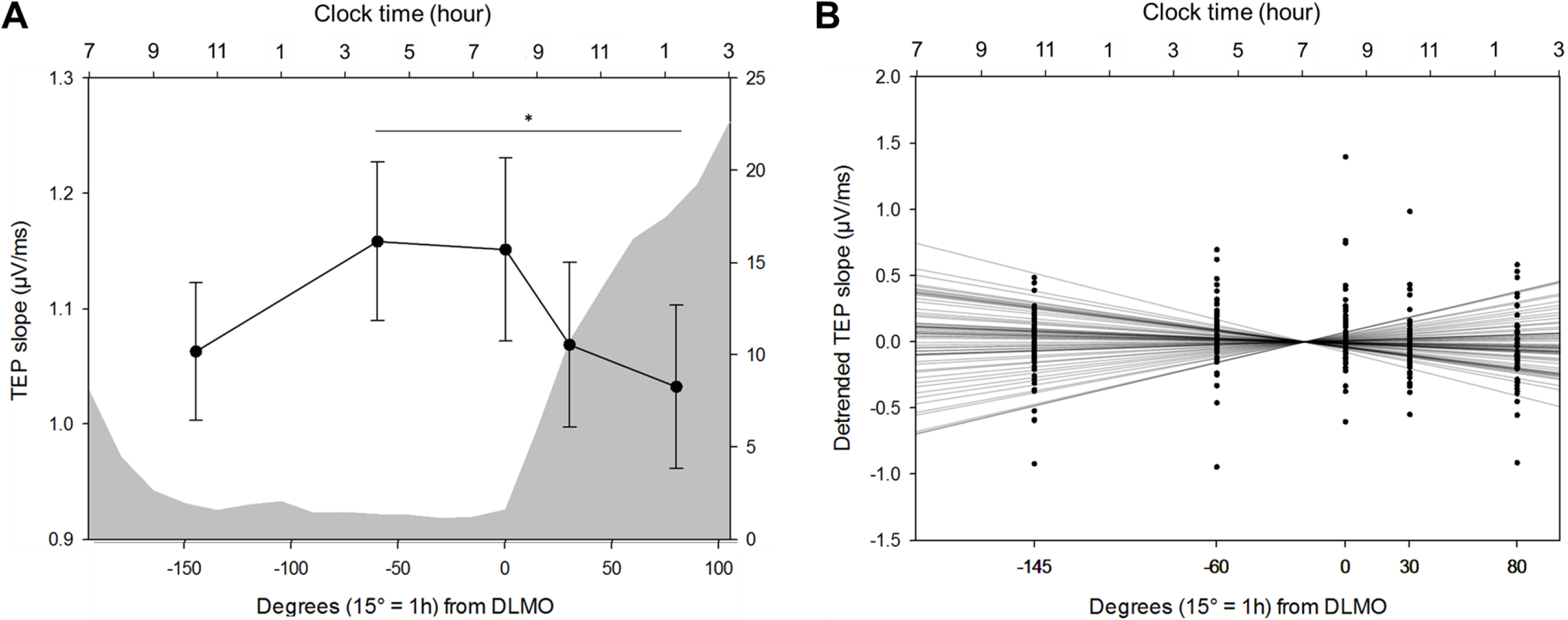
Cortical excitability dynamics as a marker of sleep-wake regulation processes. **A)** Average cortical excitability dynamics (mean ± SEM) during 20h of prolonged wakefulness over the entire sample. Grey background represents the average melatonin secretion profile (0° indicating dim light melatonin onset, i.e. the beginning of the biological night; 15° = 1h). \**p_adj_* < 0.01. **B)** Detrended cortical excitability values of all individuals and their respective linear regression lines across the 5 TMS-EEG measurements.

Whole-brain GM volume was 0.41 ± 0.04% of total intracranial volume and showed the expected reduction with increasing age (F_1,56_ = 5.86, *p* = 0.02, semi-partial *R*^2^ (*R*^2^_*β\**_) = 0.10; **Supplemental figure 2A**). Mean standardized uptake value ratio (SUVR) for Aβ and tau burden were respectively 1.16 ± 0.08 and 1.32 ± 0.10 and both showed a statistical trend in their positive association with age (Aβ: F_1,56_ = 3.74, *p* = 0.06; Tau: F_1,56_ = 3.71, p = 0.06; **Supplemental figure 2B-C**). Furthermore, whole-brain Aβ burden was strongly associated with whole-brain tau burden (F_1,55_ = 16.00, *p* = 0.0002, *R*^2^_*β\**_ = 0.23; **Supplemental figure 2D**).

### Cortical excitability dynamics during wakefulness extension

We first investigated wake-dependent changes in cortical excitability during the protocol, using a generalized linear mixed model (GLMM) including random intercept and repeated measurement autoregression [AR(1)]. Cortical excitability during prolonged wakefulness underwent significant changes with time awake after adjusting for age, sex, and education (GLMM, main effect of circadian phase; F_4,234.1_ = 4.29, *p* = 0.0023, *R*^2^_*β\**_ = 0.07; **Figure 1A**). Post-hoc analyses revealed a global decrease from the beginning to the end of the protocol, with significant differences between the second and last TMS-EEG sessions (*p*_adj_ = 0.007), and between the third and last sessions (*p*_adj_ = 0.02). Visual inspection of individual data indicated, however, an important variability in cortical excitability values and in cortical excitability dynamics. The majority of subjects (N = 35, 25 women) displayed an overall decrease in cortical excitability throughout the protocol, whereas ~40% of the sample (N = 25, 17 women) exhibited an overall increase in cortical excitability, similar to what was previously reported in young adults [24,25].

To account for this variability, we summarized cortical excitability dynamics at the individual level using a single value consisting in the regression coefficient of a linear fit across the 5 TMS-EEG measurements (**Figure 1B**). Individual residuals indicated that regression quality was good overall and regression coefficients reflected the differences between the first and last sessions of the protocol in most subjects (**Supplemental methods**). Therefore, the cortical excitability profile (CEP) obtained through the regression fit across the protocol epitomizes cortical excitability dynamics and does not reflect random signal variations. We qualified individuals with positive regression coefficient as ‘young-like’ CEP, and those with negative regression coefficient as ‘old-like’ CEP. When considered as two separate groups, young-like and old-like CEP displayed distinct temporal patterns [GLMM, group x circadian phase interaction; F_4,234.1_ = 13.69, *p* < 0.0001, *R*^2^_*β\**_ = 0.19; significant between-group post-hoc at circadian phase 30° (*p*_adj_ = 0.02) and 80° (*p* = 0.01); **Supplemental figure 1**], but did not differ in any of the demographic variables reported in **Table 1**(*p* > 0.05). Importantly, CEP was considered as a continuous variable for all statistical analyses reported below and we did not consider young-like and old-like CEP as separate groups.

### Cortical excitability dynamics during wakefulness, slow waves generation during sleep, and brain structure

We then confronted the validity of CEP as a measure of sleep-wake regulation by testing its association with slow waves generation during sleep [26]. We found that CEP was significantly related to slow wave energy (SWE), a cumulative measure of slow waves generated during NREM sleep, both in the slower range (0.75-1Hz, F_1,55_ = 5.35, *p* = 0.02, *R*^2^_*β\**_ = 0.09, **Figure 3A**) and in the higher range (1.25-4Hz, F_1,55_ = 5.47, *p* = 0.02, *R*^2^_*β\**_ = 0.09, **Figure 3B**), such that young-like CEP was associated with increased, and presumably preserved SWE. We further found that SWE in the lower 0.75-1Hz range was expectedly associated with GM volume (F_1,53_ = 7.90, *p* = 0.007, *R*^2^_*β\**_ = 0.13), as well as with whole-brain Aβ burden (F_1,53_ = 5.15, *p* = 0.03, *R*^2^_*β\**_ = 0.09, **Figure 3C**), confirming previous reports [11,27], but was not linked to whole-brain tau burden (F_1,53_ = 1.26, *p* = 0.27). In contrast, CEP was not associated with any of the brain integrity measures (GM: F_1,53_ = 0.18, *p* = 0.67; Aβ: F_1,53_ = 0.16, *p* = 0.69; Tau: F_1,53_ = 0.48, *p* = 0.49). CEP relates therefore to a gold standard measure of sleep homeostasis known to decline in aging [13,15] and to be associated with Aβ burden [11], but is not significantly linked to the hallmarks of AD neuropathology.

**Figure 3.**
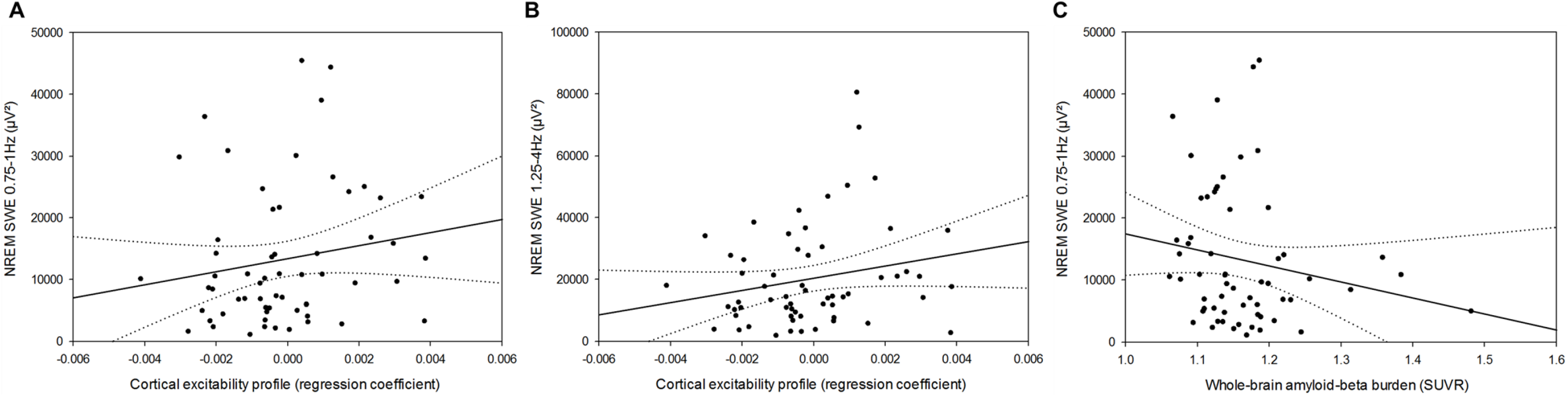
Cortical excitability, slow wave energy, and brain integrity. **A)** Positive association between CEP and cumulated frontal NREM SWE in the lower range (0.75-1Hz) during habitual sleep (F_1,55_ = 5.35, *p* = 0.02, *R^2^_β*_* = 0.09). **B)** Positive association between CEP and cumulated frontal NREM SWE in the higher range (1.25-4Hz) during habitual sleep (F_1,55_ = 5.47, *p* = 0.02, *R^2^_β*_* = 0.09). **C)** Negative association between NREM SWE (0.75-1Hz range) and whole-brain amyloid-beta burden (F_1,53_ = 5.15, *p* = 0.03, *R^2^_β*_* = 0.09). Simple regressions were used only for a visual display and do not substitute the GLMM outputs. Dotted lines represent 95% confidence interval of these simple regressions.

### Cortical excitability profile, but not slow wave energy, is associated with cognition

We tested whether CEP and NREM SWE were related to global, memory, attentional, and executive cognitive performance in GLMMs including sex, age, and education as covariates. NREM SWE was not associated with any of the cognitive measures (**Supplemental table 1**). In contrast, CEP was significantly and positively associated with global cognitive composite score, after accounting for the expected effects of age and education (F_1,55_ = 6.76, *p* = 0.01, *R*^2^_*β\**_ = 0.11, **Table 2, Figure 4A**). In other words, individuals displaying preserved cortical excitability dynamics, i.e. young-like positive CEP, had better overall cognitive performance, compared to those characterized by old-like negative CEP. Strikingly, we further found a specific and strong positive association between CEP and performance in the executive domain (F_1,55_ = 8.47, *p* = 0.005, *R*^2^_*β\**_ = 0.13, **Table 2, Figure 4B**). By contrast, neither memory (F_1,55_ = 0.41, *p* = 0.52) nor attention (F_1,55_ = 2.44, *p* = 0.12) were associated with CEP (**Table 2, Figure 4C-D**), suggesting that the link between CEP and global cognition mainly arises from the executive domain. Replacing CEP by the difference between the first and last TMS-EGG sessions led to similar statistical outcomes, ensuring that our findings are not a spurious consequence of the linear regression fit approach and thus reflect the sleep-wake-dependent regulation of the build-up of sleep need on basic brain function (**Supplemental methods; Supplemental table 2**).

**Table 2.** Associations between CEP and cognitive composite scores of global and domain-specific performance, adjusted for age, sex, and education. Statistical outputs of Generalized Linear Mixed Models with cognitive scores as dependent measures, accounting for their respective data distribution profiles. *R*^2^_*β\**_ corresponds to semi-partial *R*^2^ in GLMMs.

|  | Global performance<br>(Z-score) | Memory<br>(Z-score) | Attentional<br>(Z-score) | Executive<br>(Z-score) |
| --- | --- | --- | --- | --- |
| CEP | **$F_{1,55} = 6.76$**<br>**$p = 0.01$**<br>**$R^2_{\beta^*} = 0.11$** | $F_{1,55} = 0.41$<br>$p = 0.52$ | $F_{1,55} = 2.44$<br>$p = 0.12$ | **$F_{1,55} = 8.47$**<br>**$p = 0.005$**<br>**$R^2_{\beta^*} = 0.13$** |
| Age | **$F_{1,55} = 6.97$**<br>**$p = 0.01$**<br>**$R^2_{\beta^*} = 0.11$** | $F_{1,55} = 3.76$<br>$p = 0.06$ | **$F_{1,55} = 5.79$**<br>**$p = 0.02$**<br>**$R^2_{\beta^*} = 0.10$** | $F_{1,55} = 2.68$<br>$p = 0.11$ |
| Sex | $F_{1,55} = 0.01$<br>$p = 0.90$ | $F_{1,55} = 0.02$<br>$p = 0.90$ | $F_{1,55} = 0.05$<br>$p = 0.83$ | $F_{1,55} = 0.11$<br>$p = 0.74$ |
| Education | **$F_{1,55} = 6.18$**<br>**$p = 0.02$**<br>**$R^2_{\beta^*} = 0.10$** | $F_{1,55} = 0.14$<br>$p = 0.71$ | $F_{1,55} = 3.60$<br>$p = 0.06$ | **$F_{1,55} = 6.52$**<br>**$p = 0.01$**<br>**$R^2_{\beta^*} = 0.11$** |

**Figure 4.**
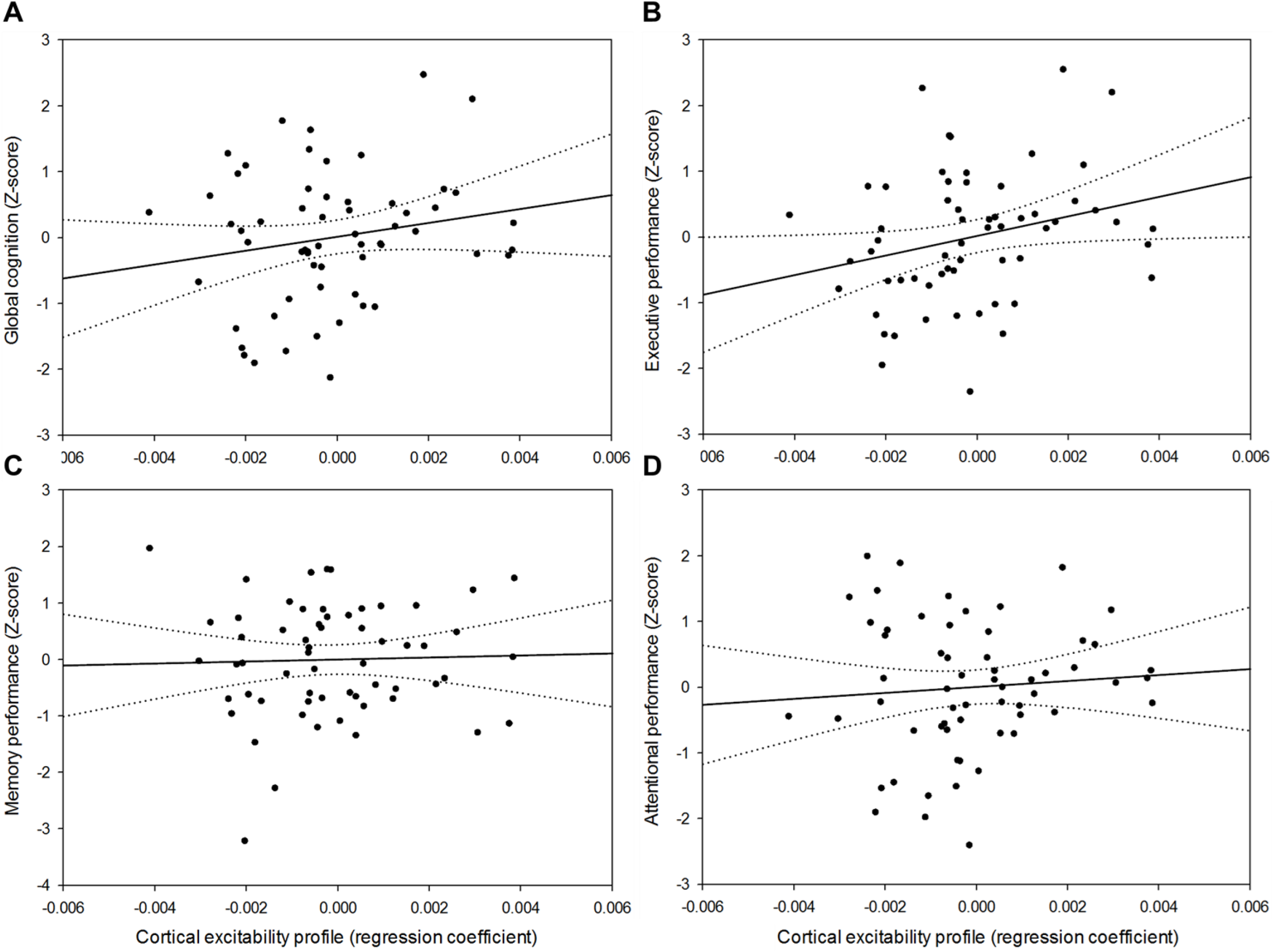
Relationships between CEP and cognition. **A)** Positive association between CEP and global cognition (F_1,55_ = 6.76, *p* = 0.01, *R^2^_β*_* = 0.11). **B)** Domain-specific positive association between CEP and performance to tasks probing executive functions (F_1,55_ = 8.47, *p* = 0.005, *R^2^_β*_* = 0.13). **C)** No significant association between CEP and memory performance (F_1,55_ = 0.39, *p* = 0.54). **D**) No significant association between CEP and attentional performance (F_1,55_ = 2.44, *p* = 0.12). Simple regressions were used only for a visual display and do not substitute the GLMM outputs. Dotted lines represent 95% confidence interval of these simple regressions.

### CEP is linked to cognition, beyond changes in brain structure associated with AD neuropathology

We found no association between the hallmarks of AD neuropathology and global, memory, attentional, and executive cognitive performance (**Table 3**). Critically, including region-specific brain integrity measures in the GLMMs seeking for associations between CEP and cognitive measures did not affect the statistical outcomes (**Table 3**); if anything, the link between CEP and executive performance became slightly stronger as suggested by semi-partial *R*^2^ values (*R*^2^_*β\**_ = 0.13 vs. *R*^2^_*β\**_ = 0.16). The associations between CEP and cognition may therefore be independent of brain integrity measures.

**Table 3.** Associations between CEP and cognitive composite scores of global and domain-specific performance, after accounting for global and region-specific brain integrity markers. Statistical outputs of Generalized Linear Mixed Models with cognitive composite scores as dependent measures, accounting for their respective data distribution profiles. When considering global cognitive performance, ‘region-specific’ Aβ and tau burden as well as GM density refer to whole-brain values. *R^2^_β*_* corresponds to semi-partial R^2^ in GLMMs.

|  | Global performance | Memory | Attentional | Executive |
| --- | --- | --- | --- | --- |
|  | (Z-score) | (Z-score) | (Z-score) | (Z-score) |
| CEP | **F<sub>1,52</sub> = 7.49**<br>**p = 0.009**<br>**R<sup>2</sup><sub>$\beta^*$</sub> = 0.13** | F<sub>1,52</sub> = 0.39<br>p = 0.54 | F<sub>1,52</sub> = 2.71<br>p = 0.11 | **F<sub>1,52</sub> = 9.71**<br>**p = 0.003**<br>**R<sup>2</sup><sub>$\beta^*$</sub> = 0.16** |
| Age | **F<sub>1,52</sub> = 7.48**<br>**p = 0.009**<br>**R<sup>2</sup><sub>$\beta^*$</sub> = 0.13** | **F<sub>1,52</sub> = 5.51**<br>**p = 0.02**<br>**R<sup>2</sup><sub>$\beta^*$</sub> = 0.10** | **F<sub>1,52</sub> = 5.22**<br>**p = 0.03**<br>**R<sup>2</sup><sub>$\beta^*$</sub> = 0.09** | F<sub>1,52</sub> = 3.16<br>p = 0.08 |
| Sex | F<sub>1,52</sub> = 0.04<br>p = 0.84 | F<sub>1,52</sub> = 0.29<br>p = 0.59 | F<sub>1,52</sub> = 0.01<br>p = 0.95 | F<sub>1,52</sub> = 0.75<br>p = 0.39 |
| Education | **F<sub>1,52</sub> = 6.47**<br>**p = 0.01**<br>**R<sup>2</sup><sub>$\beta^*$</sub> = 0.11** | F<sub>1,52</sub> = 0.99<br>p = 0.32 | F<sub>1,52</sub> = 3.22<br>p = 0.08 | **F<sub>1,52</sub> = 5.82**<br>**p = 0.02**<br>**R<sup>2</sup><sub>$\beta^*$</sub> = 0.10** |
| Region-specific GM volume | F<sub>1,52</sub> = 0.11<br>p = 0.75 | F<sub>1,52</sub> = 3.54<br>p = 0.07 | F<sub>1,52</sub> = 0.04<br>p = 0.85 | F<sub>1,52</sub> = 0.10<br>p = 0.75 |
| Region-specific Aβ burden | F<sub>1,52</sub> = 0.14<br>p = 0.71 | F<sub>1,52</sub> = 1.96<br>p = 0.17 | F<sub>1,52</sub> = 0.02<br>p = 0.88 | F<sub>1,52</sub> = 0.02<br>p = 0.90 |
| Region-specific Tau burden | F<sub>1,52</sub> = 1.17<br>p = 0.28 | F<sub>1,52</sub> = 0.66<br>p = 0.42 | F<sub>1,52</sub> = 1.07<br>p = 0.31 | F<sub>1,52</sub> = 2.51<br>p = 0.12 |

## Discussion

Healthy aging is accompanied by a disruption of sleep and wakefulness regulation that participates to age-related cognitive decline. Sleep-wake modifications are rooted in part in age-related alterations in brain integrity that can lead to AD. Previous investigations found that brain activity during sleep is related to Aβ burden, and that Aβ burden affects memory performance through modifications in brain activity during sleep [11]. This was however reported in elderly individuals (75.1 ± 3.5 years) [11], in which Aβ and tau proteins accumulation as well as neurodegeneration are likely relatively important, while daytime brain activity was not assessed. Here, we confirm that brain activity during sleep is associated with Aβ burden also in healthy late-middle aged individuals. Yet, in this relatively large (N = 60) younger sample, slow waves generation during sleep is not associated with tau burden nor with cognitive measures including memory, attention, and executive functions. We demonstrate instead that, around 60 years, sleep-wake regulation of brain activity during wakefulness is associated with cognitive fitness, independently of Aβ and tau burden and GM volume.

Previous studies reported an overall increase in cortical excitability during prolonged wakefulness in young individuals [24,25], whereas only smaller variations were detected in healthy older individuals [16]. Here, in a larger sample size of older individuals, we showed that a mild wakefulness extension results in significant changes in cortical excitability with time awake. Over the entire sample, cortical excitability globally decreased from the beginning to the end of the protocol, with marked differences between the late-afternoon and the night sessions. We further show that, amongst healthy older individuals, a significant proportion of participants displayed an overall wake-dependent increase in cortical excitability which is similar to what was previously reported in young adults [24,25]. We interpret this as a sign of preserved, or ‘young-like’, temporal dynamics of cortical excitability, which corresponds to a positive CEP or linear regression coefficient over the entire protocol. Importantly, we find that positive CEP is correlated to increased slow waves generation during habitual sleep, suggesting that individuals with ‘young-like’ CEP during wakefulness are characterized by a stronger and preserved sleep homeostasis drive during sleep [13], compared to individuals with ‘old-like’ CEP. This strongly suggests that CEP is a measure of the active brain during wakefulness that reflects individuals’ preservation of sleep-wake regulation processes.

Critically, we found that higher CEP is associated with better overall cognitive performance, demonstrating a significant relationship between cortical excitability dynamics during wakefulness and cognition, which are both measured in an active and awake brain. In-depth cognitive phenotyping showed that this relationship was mainly driven by the performance in the executive domain. Executive functions refer to high-order cognitive processes (flexibility, inhibition, updating, etc.) needed for behavioral adjustment according to ongoing goals when facing new or complex situations [28]. Variations in executive performance assessed during sleep deprivation were previously found to be associated with cortical excitability dynamics in older and younger individuals [16]. Here, we further show that cognitive ability, as an individual trait measured outside a sleep deprivation protocol, is significantly associated with sleep-wake regulation of basic brain function. Executive functions influence other cognitive domains and are often seen as central in age-related cognitive decline to remain adapted to the environment and sustain day-to-day functioning in complete autonomy [29]. In addition, executive functions are considered to depend mainly on the frontal cortex processes, and their underlying cortical networks undergo significant changes in healthy aging [30]. These results therefore suggest that preserved frontal CEP may constitute a marker of cognitive fitness in aging, and particularly so in the executive domain.

Despite the relative youth of our sample (mean 59.6 ± 5.5 years), the absence of association between cognitive measures and brain integrity markers of GM volume and protein burdens may appear surprising. Longitudinal studies have shown, however, that the association between some aspects of cognition and brain structure was especially apparent in participants aged 65 years and over [31]. Subtle differences in cognitive performance in relation to proteins accumulation or GM reduction may therefore not appear in our sample. This may also underlie the absence of significant association between sleep slow waves generation and tau burden that has been recently reported in an older sample of healthy individuals (73.8 ± 5.3 years) [10]. Furthermore, our sample is biased towards individuals with higher education (mean 15.5 ± 3.2 years) with cognitive and brain reserves that may compensate for early alteration of brain integrity [3,32].. Alternatively, the composite scores for each cognitive domain may not be sensitive enough to be related to these early brain alterations. Nonetheless, our findings indicate that sleep-wake regulation of brain activity during wakefulness, as measured by the dynamics of cortical excitability during a mild wakefulness extension, is either more sensitive than brain integrity markers to isolate associations with cognition in aging, or sensitive to aspects of cognition that undergo influences distinct from protein accumulations and neurodegeneration. Another explanation to our results might involve the soluble forms of Aβ and tau, as oligomers of both proteins were shown to alter neuronal function [33,34]. Currently these oligomers cannot be reliably measured in vivo and could not be accounted for in this experiment.

Furthermore, age-related molecular changes potentially underlying sleep need have been reported [35]. These may influence the local or global genetic and molecular machineries underlying circadian rhythmicity and sleep homeostasis, and in turn affect cortical excitability. Age-related modifications modulate the impact of the basal forebrain, subcortical and brainstem ascending activating system on global brain activity [36]. In silico modeling of wake-dependent cortical excitability changes in young adults suggests that the fluctuations in the balance between excitation and inhibition within cortical networks may affect the observed variations during prolonged wakefulness [37]. However, this remains unexplored in healthy older individuals. It might also be the case that changes in cortical function stems from AD-related alteration of subcortical structures. In addition to the locus coeruleus, neurodegeneration of the suprachiasmatic nuclei (SCN; site of the master circadian clock) has been reported in AD [38] while SCN network uncoupling is found in normal aging [39]. Future investigations should also examine whether frontal cortical excitability dynamics in particular are related to cognitive changes in aging when compared to other parts of the brain and to other aspects of brain functions affected by the interplay between circadian rhythmicity and sleep homeostasis. Furthermore, the predictive value of CEP assessment for subsequent cognitive decline and risk of developing dementia remains to be investigated in a longitudinal protocol.

This study presents several strengths. The use of TMS-EEG allows for a direct measure of cortical responsiveness while bypassing sensory systems and it mimics active brain processing without confounding biases. The prolonged wakefulness protocol is performed under strictly controlled constant routine conditions to control for multiple factors that could affect wakefulness and sleep parameters, such as light exposure or physical activity [14]. In addition, we performed a comprehensive multi-modal assessment of brain structure, including two PET scans and MRI for the hallmarks of AD pathophysiology, as well as an extensive neuropsychological investigation. Furthermore, sleep-wake history was controlled prior to wake extension and exclusion criteria ensured most risk factor favoring cognitive decline were not present in the sample (e.g. diabetes, smoking, alcohol abuse, depression, etc.) [40]. Finally, the relative young age of the participants reduces the accumulation of minor health issues associated with advanced age (e.g. hyper-tension, diabetes, overweight, etc.), which can inherently affect findings in samples of elderly individuals. This research also has several limitations, however. Its cross sectional nature does not allow us to comment on the future cognitive trajectory of participants. The reported effect sizes show that CEP does explain but a small part of variance, suggesting that its link with cognitive fitness is modest. This was expected given the numerous factors that affect cognitive trajectories [40] and the relatively young age and good overall health status of our sample. It is, in fact, quite remarkable that we were nonetheless able to isolate a link between sleep-wake regulation of brain activity and cognition in such sample, suggesting that it may be a very important link to successful cognitive aging. While the contribution of the homeostatic process is most obvious in our data, a longer protocol covering the whole circadian cycle would help disentangle the respective modulation of CEP by the circadian system [14]. In addition, ^[18F]^THK-5351 presents some unspecific binding, particularly around the fornix and basal ganglia [41]. We took this into account by excluding these portions of the brain from all tau burden measures. The observed links between ^[18F]^THK-5351 uptake and both age and Aβ burden strongly support that our measure of tau burden, although potentially imperfect, was meaningful. Finally, we did not consider other age-related changes of brain integrity, such as cerebrovascular pathology, which are extremely common (up to 50%) as a mixed pathology in individuals with Alzheimer’s dementia [42], and Lewy bodies pathology, which shares some genetic risk with Alzheimer’s disease [43].

Aging is the ultimate challenge that the brain has to face in order to maintain optimal cognition across the lifespan. The bidirectional detrimental interaction between disturbed sleep-wake regulation and AD pathogenesis suggest that sleep-wake interventions could be promising means to reduce the risk of dementia [6]. Here, we provide compelling evidence that sleep-wake regulation influences cognition in healthy older individuals, particularly in the executive domain, and beyond the changes in brain integrity that can ultimately lead to dementia. Since both sleep homeostasis and circadian rhythmicity show significant alteration as early as age 40 [13,15], our results further reinforces the idea that sleep and wakefulness could be acted upon to improve individual cognitive health trajectory early in the lifespan. Our findings could have therefore implications for the understanding of brain mechanisms underlying the maintenance of cognitive health in normal and pathological aging, and for potential early intervention targets.

## Methods

### Study design and participants

Between June 15, 2016, and July 28, 2018, healthy older individuals aged 50-70 years were enrolled for this multi-modal cross-sectional study after giving their written informed consent, and received a financial compensation. This research was approved by the Ethics Committee of the Faculty of Medicine at the University of Liège, Belgium. Exclusion criteria were: clinical symptoms of cognitive impairment (Dementia rating scale < 130; Mini mental state examination < 27); Body Mass Index (BMI) ≤ 18 and ≥ 29; recent psychiatric history or severe brain trauma; addiction, chronic medication affecting the central nervous system; smoking, excessive alcohol (> 14 units/week) or caffeine (> 5 cups/day) consumption; shift work in the past 6 months; transmeridian travel in the past two months; anxiety, as measured by the 21-item self-rated Beck Anxiety Inventory (BAI ≥ 10); depression, as assessed by the 21-item self-rated Beck Depression Inventory (BDI ≥ 14). Participants with sleep apnea (apnea-hypopnea index ≥ 15/hour) were excluded based on an in-lab screening night of polysomnography. One participant was excluded from the sample for all analyses because of outlier values on both PET assessments (> 6 standard deviations from the mean). Demographic characteristics of the final study sample are described in **Table 1**.

### Magnetic resonance imaging

High-resolution structural MRI was performed on a 3-Tesla MR scanner (MAGNETOM Prisma, Siemens). For each participant, multi-parameter mapping volumes (*i.e*. T1-weighted, Proton Density (PD)-weighted, Magnetization Transfer (MT)-weighted) were acquired. We estimated individuals’ total intracranial volume and whole-brain GM volume based on the MT-weighted image, using the SPM12 toolbox (https://www.fil.ion.ucl.ac.uk/spm/). For regional quantification of GM, volumes of interests (VOIs) were first determined using the Automated Anatomical Labeling atlas (AAL2) [44]. MT-weighted images were spatially normalized into a study-specific template with the Diffeomorphic Anatomical Registration Through Exponentiated Algebra (DARTEL) toolbox [45]. VOIs were then applied on segmented normalized MT-weighted images and combined to extract GM volume in brain regions underlying each cognitive domain (**Supplemental table 3**).

### PET imaging

Aβ-PET imaging was performed with ^[18F]^Flutemetamol, and tau-PET imaging was done with ^[18F]^THK-5351. For both radiotracers, all participants received a single dose of their respective radioligands in an antecubital vein (target dose 185 MBq). Aβ-PET image acquisitions started 85 minutes after injection, and 4 frames of 5 minutes were obtained, followed by a 10-minute transmission scan. For tau-PET, transmission scan was acquired first and dynamic image acquisitions started immediately after injection, consisting in 32 frames (with increasing time duration). All PET images were reconstructed using filtered back-projection algorithm including corrections for measured attenuation, dead time, random events, and scatter using standard software (ECAT 7.1, Siemens/CTI, Knoxville, TN). Motion correction was performed using automated realignment of frames without reslicing. A PET sum image was created using all frames for Aβ-PET, and using the 4 frames corresponding to the time window between 40 and 60 min for tau-PET. PET sum images were reoriented manually according to MT-weighted structural MRI volume and coregistered to structural MRI using the MT-weighted volume. PET sum images were further corrected for partial volume effect (PETPVC toolbox, iterative Yang method [46]) and spatially normalized using the MRI study-specific template. Standardized uptake value ratio (SUVR) was calculated using the cerebellum GM as the reference region. The VOIs used for GM analysis were applied to normalized PET sum images to estimate regional SUVR of each radiotracer in cognitive domain-specific regions. ^[18F]^THK-5351 radiotracer shows some unspecificity for tau, particularly over the basal ganglia, which was taken into account by excluding basal ganglia from all computations of PET SUVR values for tau-PET.

### Cognitive assessment

Upon arrival for the wake-extension protocol and prior to being placed in dim-light (~7.5 hours before habitual bedtime), participants were administered the first part (~1h) of the extensive neuropsychological assessment including: 1) Mnemonic Similarity Task (MST); 2) Category Verbal Fluency (“letter” and “animals”); 3) Digit Symbol Substitution Test; 4) Visual N-Back (1-, 2-*, and 3-back variants); and 5) Choice Reaction Time. On another day while well-rested and during the day (from 12 to 6 hours before habitual bedtime), the second part of the neuropsychological assessment was administered. This ~1.5h session included: 1) Direct and Inverse Digit Span; 2) Free and Cued Selective Reminding Test; 3) Stroop Test; 4) Trail Making Test (TMT part A and B); and 5) D2 Attention Test. The memory function composite score included FCSRT (sum of all free recalls) and MST (recognition memory score). The executive function composite score comprised verbal fluency tests (2-min score for letter and animal variants), the digit span (inverse order), TMT (part B), N-Back (3-back variant), and Stroop (number of errors for interfering items). The attentional function composite score included DSST (2-min score), TMT (part A), N-Back (1-back variant), D2 (Gz-F score), and CRT (reaction time to dissimilar items). We computed a composite score for each cognitive domain based on the sum of Z-scores on domain-related tasks, with higher scores reflecting better performance. Composite score for global cognitive performance score consisted of the standardized sum of the domain-specific composite scores.

### Sleep assessment and spectral power analysis

For 7 days prior to the wake-extension protocol, participants followed a regular sleep-wake schedule (± 30 min), in agreement with their preferred bed and wake-up times. Compliance was verified using sleep diaries and wrist actigraphy (Actiwatch^©^, Cambridge Neurotechnology, UK). The day before the wake-extension protocol, participants arrived to the laboratory 8 hours before their habitual bedtime and were kept in dim light (< 5 lux) for 6.5 hours preceding bedtime. Their habitual sleep was then recorded in complete darkness under EEG (baseline night, **Figure 1A**). Baseline night data were acquired using N7000 amplifiers (EMBLA). The electrode montage consisted of 11 EEG channels (F3, Fz, F4, C3, Cz, C4, P3, Pz, P4, O1, O2), 2 bipolar EOGs, and 2 bipolar EMGs. Scoring of baseline night in 30-s epochs was performed automatically using a validated algorithm (ASEEGA, PHYSIP, Paris, France) [47]. An automatic artefact detection algorithm with adapting thresholds [48] was further applied on scored data. Power spectrum was computed for each channel using a Fourier transform on successive 4-s bins, overlapping by 2-s., resulting in a 0.25 Hz frequency resolution. The night was divided into 30 min periods, from sleep onset until lights on. For each 30 min period, slow wave energy (SWE) was computed as the sum of generated power in the delta band, both for the lower range (0.75-1Hz) and higher range (1.25-4Hz), during all the NREM 2 (N2) and NREM 3 (N3) epochs of the given period, after adjusting for the number of N2 and N3 epochs to account for artefacted data. As the frontal regions are most sensitive to sleep-wake history [13], SWE was considered over the frontal electrodes (F3, Fz, F4).

### Wake-extension protocol

The wake-extension protocol followed the baseline night and consisted of 20h of continuous wakefulness under strictly controlled constant routine conditions, *i.e.* in-bed semi-recumbent position (except for scheduled bathroom visits), dim light < 5 lux, temperature ~19°C, regular isocaloric food intake, no time-of-day information, and sound-proofed rooms. The protocol schedule was adapted to individual sleep-wake time, and lasted up to the theoretical mid-sleep time (e.g. ca. 07:00 AM-03:00 AM) to create a moderate wakefulness extension challenge. Hourly saliva samples were collected for subsequent melatonin assays, allowing *a posteriori* data realignment and interpolation based on individual endogenous circadian timing. Cortical excitability over the frontal cortex was measured 5 times throughout the protocol, using TMS-EEG, with increased frequency around the circadian wake-maintenance zone, as it represents a critical period around which the interplay between sleep homeostasis and the circadian system show important changes.

### TMS-EEG assessment

One TMS-EEG session was performed prior to the wake-extension protocol to determine optimal stimulation parameters (i.e. location, orientation, and intensity) that allowed for EEG recordings free of muscular and magnetic artefacts. As in previous experiments [16,24,25], the target location was in the superior frontal gyrus. For all TMS-EEG recordings, pulses were generated by a Focal Bipulse 8-Coil (Nexstim, Helsinki, Finland). Interstimulus intervals were randomized between 1900 and 2200 ms. TMS-evoked responses were recorded with a 60-channel TMS-compatible EEG amplifier (Eximia, Helsinki, Finland), equipped with a proprietary sample-and-hold circuit which provides TMS artefact free data from 5 ms post stimulation. Electrooculogram (EOG) was recorded with two additional bipolar electrodes. EEG signal was band-pass filtered between 0.1 and 500 Hz and sampled at 1450 Hz. Before each recording session, electrodes impedance was set below 5 kΩ. Each TMS-EEG session included ~250 trials (mean = 252 ± 15). Auditory EEG potentials evoked by the TMS clicks and bone conductance were minimized by diffusing a continuous white noise through earphones and applying a thin foam layer between the EEG cap and the TMS coil. A sham session, consisting in 30 TMS pulses delivered parallel to the scalp with noise masking, was administered to verify the absence of auditory EEG potentials. TMS-EEG data were preprocessed as previously described [16]. Cortical excitability was computed as the slope at the inflexion point of the first component of the TMS-evoked EEG potential on the electrode closest to the stimulation hotspot. For each participant, CEP value was defined as the regression coefficient of a linear fit (first order ‘polyfit’ MATLAB function) across the 5 TMS-EEG measurements at the individual level.

### Melatonin assessment

Salivary melatonin was measured by radioimmunoassay. The detection limit of the assay for melatonin was 0.8 ± 0.2 pg/l using 500 μl volumes. Dim light melatonin onset (DLMO) times were computed for each participant using the hockey-stick method, with ascending level set to 2.3 pg/ml (Hockey-Stick software v1.5) [49]. The circadian phase of all TMS-EEG data points was estimated relative to individual DLMO time (*i.e.* phase 0°; 15° = 1h). Based on this, cortical excitability measures were resampled following linear interpolation at the theoretical phases of the TMS-EEG sessions of the protocol (−145°, −60°, 0°, 30°, 80°).

### Statistics

Statistical analyses were performed using Generalized Linear Mixed Models (GLMMs) in SAS 9.4 (SAS Institute, Cary, NC). Dependent variables distribution was first determined using “allfitdist” function in MATLAB and GLMMs were adjusted accordingly. All statistical models were adjusted for age, sex and education. Statistical significance was set at *p* < 0.05. Simple regressions were used for visual display only and not as a substitute of the full GLMM statistics. Degrees of freedom were estimated using Kenward-Roger’s correction. *P*-values in post-hoc contrasts (difference of least square means) were adjusted for multiple testing with Tukey’s procedure. Semi-partial *R*^2^ (*R*^2^_*β\**_) values were computed to estimate effect size of significant fixed effects in all GLMMs [50].

## Author contributions

Study concept and design: E.S., P.M., C.P., C.B., F.C., and G.V. Data acquisition, analysis and interpretation: all authors. Administrative, technical, or material support: E.S., A.L., P.M., and C.P. Drafting of the manuscript: MV, DC, and GV. Manuscript revising: all authors.

## Acknowledgments

We thank M. Blanpain, M. Cerasuolo, E. Lambot, C. Hagelstein, S. Laloux, E. Balteau, A. Claes, C. Degueldre, B. Herbillon, P. Hawotte, and B. Lauricella for their help in different steps of the study.

## Data and materials availability

The authors declare that the data supporting the findings of this study are available from the corresponding author upon request.

