## Supplemental methods for "Preserved wake-dependent cortical excitability dynamics predict cognitive fitness beyond age-related brain alterations"

### Cortical excitability profile index validation

When analysing the time course of cortical excitability values across the 5 phases of TMS-EEG acquisitions, residuals from the linear modelling verified the goodness-of-fit across the sample. Furthermore, we computed the difference between the last and the first TMS-EEG sessions (TEP slope S5-S1), so that the ‘young-like’ increase in cortical excitability during the biological night compared to the biological day would be also reflected in a positive value. The proportions of young-like vs. old-like subjects measured with CEP or TEP slope S5-S1 were identical (*i.e.* 25 young-like vs. 35 old-like in both cases), and the individual attribution to young-like or old-like profiles were highly matched between the two indices (Phi coefficient = 0.86; only 4 individuals were not attributed to the same group).

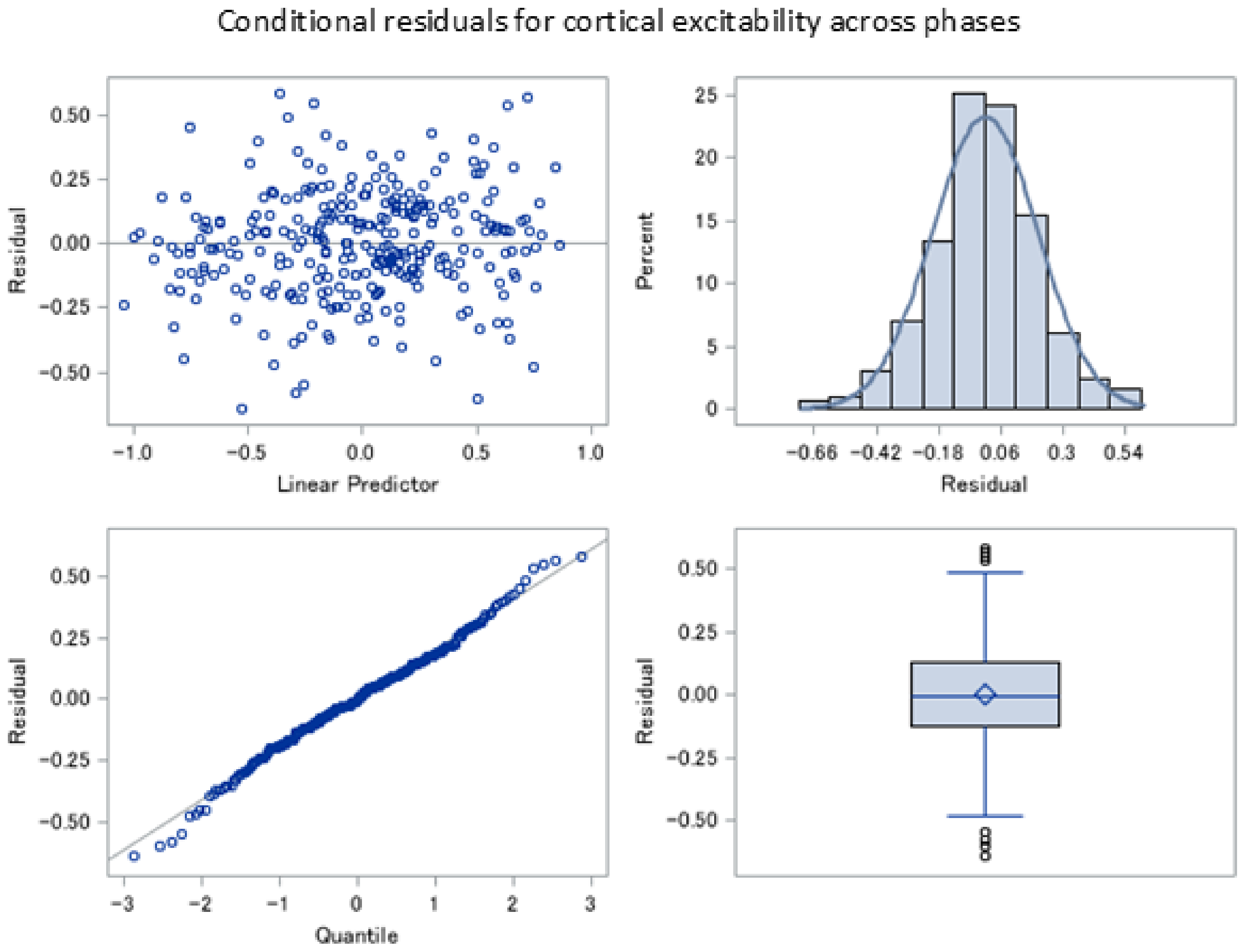

**Supplemental figures and tables**

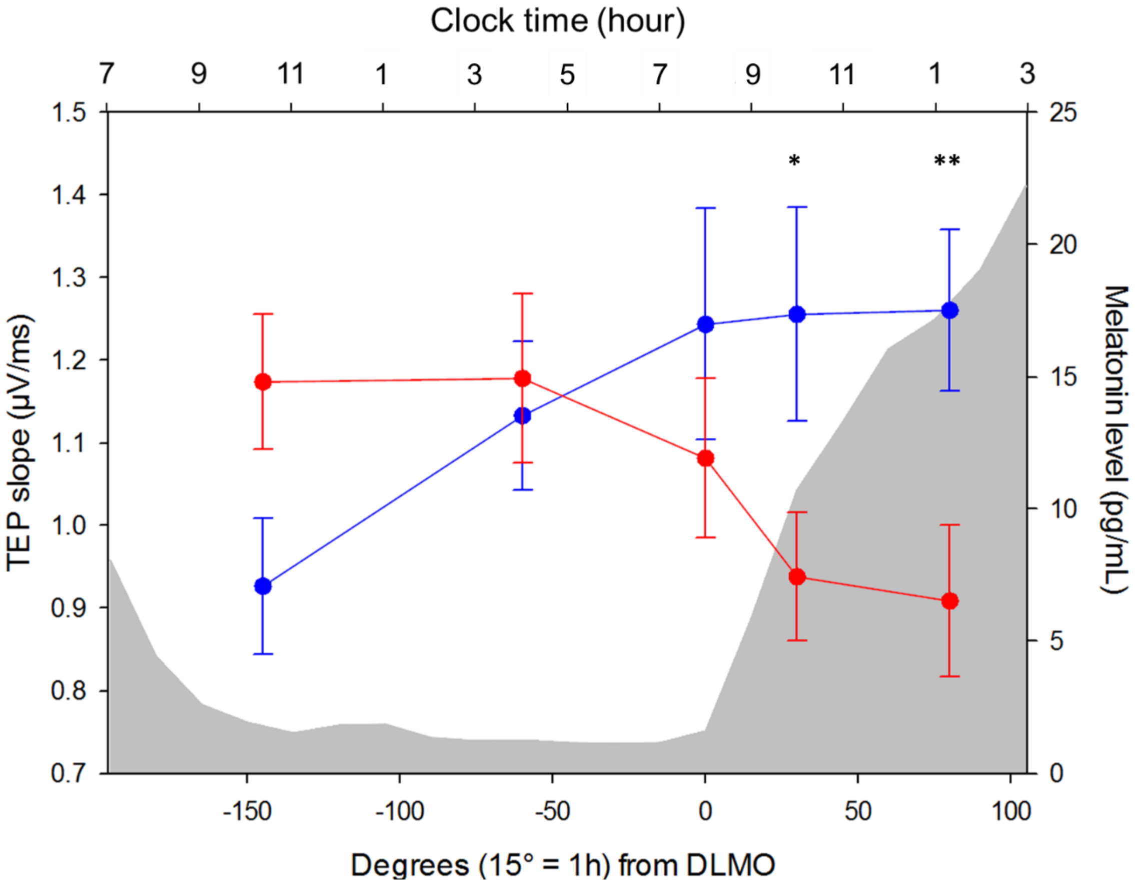


**Supplemental figure 1.** Averaged cortical excitability dynamics of individuals displaying young-like cortical excitability profile (CEP) (blue, n = 25) vs. individuals who display old-like CEP (red, n = 35). * $p_{adj} < 0.05$; ** $p_{adj} < 0.01$. Importantly, except for analyses related to this figure, CEP was considered as a continuous variable for all statistical analyses in the present study.

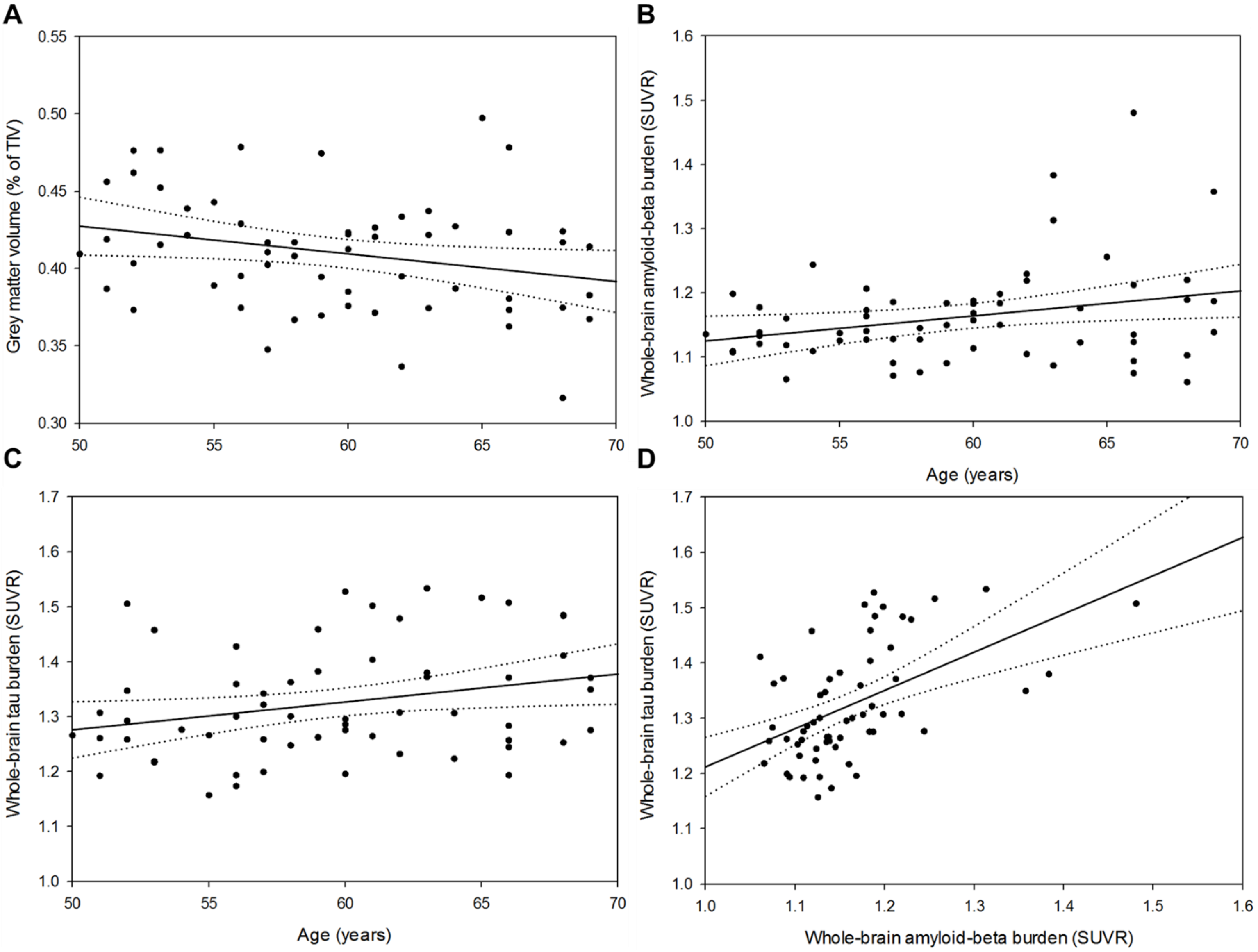


**Supplemental figure 2. Relationships between age and brain integrity measures. A)** Negative association between age and whole-brain GM volume ($F_{1,56}$ = 5.86, $p$ = 0.02, $R^2_{\beta*}$ = 0.10). **B)** Statistical trend towards a positive association between age and whole-brain Aβ burden ($F_{1,56}$ = 3.74, $p$ = 0.06). **C)** Statistical trend towards a positive association between age and whole-brain tau burden ($F_{1,56}$ = 3.71, $p$ = 0.06). **D)** Positive association between whole-brain Aβ protein burden and whole-brain tau protein burden ($F_{1,55}$ = 16.00, $p$ = 0.0002, $R^2_{\beta*}$ = 0.23). Simple regressions were used only for a visual display of the direction of the associations. Dotted lines represent 95% confidence interval of these simple regressions. The narrow age range of our relatively young sample most likely explains that we observe trends rather than significant links between age and Aβ or tau burdens. Nonetheless, the fact that both PET markers show the expected positive association with increasing age support their validity as markers of brain integrity.

**Supplemental table 1. Associations between NREM SWE (0.75-4Hz range*) and cognition.**

Associations between NREM SWE (0.75-4Hz range*) and cognitive composite scores of global and domain-specific performance.

| | Global performance (Z-score) | Memory (Z-score) | Attentional (Z-score) | Executive (Z-score) |
|---|---|---|---|---|
| NREM SWE (0.75-4Hz) | $F_{1,55}$ = 0.42 | $F_{1,55}$ = 1.42 | $F_{1,55}$ = 0.18 | $F_{1,55}$ = 1.74 |
| | *p* = 0.52 | *p* = 0.24 | *p* = 0.67 | *p* = 0.19 |
| Age | $F_{1,55}$ = 2.72 | **$F_{1,55}$ = 4.40** | $F_{1,55}$ = 3.34 | $F_{1,55}$ = 0.19 |
| | *p* = 0.11 | ***p* = 0.04** | *p* = 0.07 | *p* = 0.67 |
| | | **$R^2_{\beta*}$ = 0.07** | | |
| Sex | $F_{1,55}$ = 0.02 | $F_{1,55}$ = 0.29 | $F_{1,55}$ = 0.13 | $F_{1,55}$ = 0.04 |
| | *p* = 0.89 | *p* = 0.59 | *p* = 0.72 | *p* = 0.85 |
| Education | **$F_{1,55}$ = 4.59** | $F_{1,55}$ = 0.08 | $F_{1,55}$ = 2.99 | **$F_{1,55}$ = 4.81** |
| | ***p* = 0.04** | *p* = 0.78 | *p* = 0.09 | ***p* = 0.03** |
| | **$R^2_{\beta*}$ = 0.08** | | | **$R^2_{\beta*}$ = 0.08** |

Associations between NREM SWE (0.75-4Hz range*) and cognitive composite scores of global and domain-specific performance, accounting for global and region-specific brain integrity markers.

| | Global performance (Z-score) | Memory (Z-score) | Attentional (Z-score) | Executive (Z-score) |
|---|---|---|---|---|
| NREM SWE (0.75-4Hz) | $F_{1,52}$ = 0.49 | $F_{1,52}$ = 0.20 | $F_{1,52}$ = 0.14 | $F_{1,52}$ = 1.29 |
| | *p* = 0.49 | *p* = 0.66 | *p* = 0.71 | *p* = 0.26 |
| Age | $F_{1,52}$ = 3.36 | **$F_{1,52}$ = 5.26** | $F_{1,52}$ = 3.23 | $F_{1,52}$ = 0.59 |
| | *p* = 0.07 | ***p* = 0.03** | *p* = 0.08 | *p* = 0.45 |
| | | **$R^2_{\beta*}$ = 0.09** | | |
| Sex | $F_{1,52}$ = 0.03 | $F_{1,52}$ = 0.08 | $F_{1,52}$ = 0.06 | $F_{1,52}$ = 0.02 |
| | *p* = 0.87 | *p* = 0.78 | *p* = 0.82 | *p* = 0.88 |
| Education | **$F_{1,52}$ = 4.72** | $F_{1,52}$ = 0.71 | $F_{1,52}$ = 2.57 | **$F_{1,52}$ = 4.52** |
| | ***p* = 0.03** | *p* = 0.40 | *p* = 0.12 | ***p* = 0.04** |
| | **$R^2_{\beta*}$ = 0.08** | | | **$R^2_{\beta*}$ = 0.08** |
| Region-specific GM volume | $F_{1,52}$ = 0.17 | $F_{1,52}$ = 2.41 | $F_{1,52}$ = 0.03 | $F_{1,52}$ = 0.01 |
| | *p* = 0.68 | *p* = 0.13 | *p* = 0.85 | *p* = 0.92 |
| Region-specific Aβ burden | $F_{1,52}$ = 0.14 | $F_{1,52}$ = 1.66 | $F_{1,52}$ = 0.01 | $F_{1,52}$ = 0.11 |
| | *p* = 0.71 | *p* = 0.20 | *p* = 0.91 | *p* = 0.74 |
| Region-specific Tau burden | $F_{1,52}$ = 0.53 | $F_{1,52}$ = 0.79 | $F_{1,52}$ = 0.67 | $F_{1,52}$ = 1.14 |
| | *p* = 0.47 | *p* = 0.38 | *p* = 0.42 | *p* = 0.29 |

* For simplicity, results are presented for the entire range of SWE, *i.e.* lower (0.75-1Hz) and higher (1.25-4Hz) ranges combined, but the statistical outcomes are the same when considering both ranges separately.

**Supplemental table 2. Associations between the difference in cortical excitability values between the last and the first TMS-EEG sessions (S5–S1) and cognitive composite scores of global and domain-specific performance**, adjusted for age, sex, and education. Statistical outputs of Generalized Linear Mixed Models with cognitive scores as dependent measures, accounting for their respective data distribution profiles. $R^2_{\beta*}$ correspond to semi-partial $R^2$ in GLMMs.

| | Global performance (Z-score) | Memory (Z-score) | Attentional (Z-score) | Executive (Z-score) |
|---|---|---|---|---|
| TEP slope (S5-S1) | $F_{1,55}$ = 2.90<br>$p$ = 0.09 | $F_{1,55}$ = 0.10<br>$p$ = 0.76 | $F_{1,55}$ = 1.31<br>$p$ = 0.26 | **$F_{1,55}$ = 4.75**<br>**$p$ = 0.03**<br>**$R^2_{\beta*}$ = 0.08** |
| Age | **$F_{1,55}$ = 5.27**<br>**$p$ = 0.03**<br>**$R^2_{\beta*}$ = 0.10** | $F_{1,55}$ = 2.85<br>$p$ = 0.10 | **$F_{1,55}$ = 5.09**<br>**$p$ = 0.03**<br>**$R^2_{\beta*}$ = 0.08** | $F_{1,55}$ = 1.83<br>$p$ = 0.18 |
| Sex | $F_{1,55}$ = 0.04<br>$p$ = 0.85 | $F_{1,55}$ = 0.01<br>$p$ = 0.92 | $F_{1,55}$ = 0.03<br>$p$ = 0.87 | $F_{1,55}$ = 0.18<br>$p$ = 0.67 |
| Education | **$F_{1,55}$ = 5.16**<br>**$p$ = 0.03**<br>**$R^2_{\beta*}$ = 0.09** | $F_{1,55}$ = 0.09<br>$p$ = 0.77 | $F_{1,55}$ = 3.25<br>$p$ = 0.08 | **$F_{1,55}$ = 5.49**<br>**$p$ = 0.02**<br>**$R^2_{\beta*}$ = 0.09** |

**Supplemental table 3. Masks of domain-specific brain regions computed with Automated Anatomical Labeling (*AAL2*) atlas.** The table lists the anatomical brain regions used to create the mask for the executive function [1,2], memory function [3,4], and attentional function [5,6]. Names in *italic* refer to the labels of the regions in *AAL2* atlas (http://www.gin.cnrs.fr/en/tools/aal-aal2/).

| Memory function | Attentional function | Executive function |
|---|---|---|
| Medial superior frontal<br>(*Frontal_Sup_Medial*) | Superior frontal<br>(*Frontal_Sup_2*) | Superior frontal<br>(*Frontal_Sup_2*) |
| Medial orbitofrontal<br>(*Frontal_Med_Orb*) | Middle frontal<br>(*Frontal_Mid_2*) | Medial superior frontal<br>(*Frontal_Sup_Medial*) |
| Posterior cingulate<br>(*Cingulate_Post*) | Inferior frontal, pars triangularis<br>(*Frontal_Inf_Tri*) | Middle frontal<br>(*Frontal_Mid_2*) |
| Hippocampus<br>(*Hippocampus*) | Superior parietal<br>(*Parietal_Sup*) | Inferior frontal, pars opercularis<br>(*Frontal_Inf_Oper*) |
| Parahippocampal<br>(*ParaHippocampal*) | Inferior parietal<br>(*Parietal_Inf*) | Inferior frontal, pars triangularis<br>(*Frontal_Inf_Tri*) |
| Angular<br>(*Angular*) | Anterior cingulate<br>(*Cingulate_Ant*) | Inferior frontal, pars orbitalis<br>(*Frontal_Inf_Orb_2*) |
| Precuneus<br>(*Precuneus*) | Superior temporal<br>(*Temporal_Sup*) | Medial orbitofrontal<br>(*Frontal_Med_Orb*) |
| | Thalamus<br>(*Thalamus*) | Anterior cingulate<br>(*Cingulate_Ant*) |
| | Fusiform<br>(*Fusiform*) | Superior parietal<br>(*Parietal_Sup*) |
| | Precentral<br>(*Precentral*) | Inferior parietal<br>(*Parietal_Inf*) |
| | Postcentral<br>(*Postcentral*) | |
